# Urbanization and market-integration have strong, non-linear effects on cardio-metabolic health in the Turkana

**DOI:** 10.1101/756866

**Authors:** Amanda J. Lea, Dino Martins, Joseph Kamau, Michael Gurven, Julien F. Ayroles

**Affiliations:** Department of Ecology and Evolution, Princeton University, Princeton, New Jersey, USA; Lewis Sigler Institute for Integrative Genomics, Princeton University, Princeton, New Jersey, USA; Mpala Research Centre, Nanyuki, Kenya; Department of Biochemistry, School of Medicine, University of Nairobi, Nairobi, Kenya; Institute of Primate Research, National Museums of Kenya, Nairobi, Kenya; Department of Anthropology, University of California: Santa Barbara, Santa Barbara, California, USA

**Keywords:** pastoralism, metabolic health, industrialization, market integration, evolutionary mismatch, Turkana

## Abstract

Cardio-metabolic disease is a leading cause of death worldwide, with high prevalence in western, industrialized societies relative to developing nations and subsistence-level populations. This stark difference has been attributed to the dietary and lifestyle changes associated with industrialization, but current work has relied on health comparisons between separate, genetically distinct populations to draw conclusions. To more robustly determine how lifestyle impacts health, we collected interview and health biomarker data from a single population undergoing a rapid lifestyle transition. Specifically, we sampled Turkana individuals who practice subsistence-level, nomadic pastoralism (the traditional, ancestral way of life for this group), as well as individuals who no longer practice pastoralism and engage either minimally or strongly with the market economy. Comparisons across this lifestyle gradient revealed clear, non-linear effects of industrialization: only individuals with highly urban, market-integrated lifestyles experience increases in BMI, body fat, blood pressure, and other biomarkers of cardio-metabolic health. These health differences are partially mediated by increased consumption of refined carbohydrates, and more strongly by fine-scale measures of urbanicity. Finally, because many Turkana are transitioning between rural and urban areas within their lifetime, we were able to show that being born in an urban area is associated with worse adult metabolic health, independent of adult lifestyle. Together, these analyses provide comprehensive insight into the timing, magnitude, and causes of health declines in urban, industrialized groups – an area of critical study given the massive public health burden of cardio-metabolic disease and the rate at which developing nations are experiencing lifestyle transitions.

**SIGNIFICANCE:** The “mismatch” between evolved human physiology and western, industrialized lifestyles is thought to explain to the current epidemic of cardiovascular disease (CVD). However, this hypothesis has been difficult to test in real time. To do so, we studied a traditional pastoralist group—the Turkana—that is currently transitioning from their ancestral way of life to an urban, industrialized lifestyle. We found that Turkana who move to cities exhibit poor cardio-metabolic health, partially because of a shift toward “western diets” high in carbohydrates. We also show that early life urbanicity independently predicts adult health, such that life-long city dwellers will experience the greatest CVD risk. Our work thus uncovers the timing, magnitude, and evolutionary causes of a major health gradient.

## INTRODUCTION

Over the last several decades, it has become increasingly clear that the spread of ‘western’, industrialized lifestyles is contributing to a rapid, worldwide rise in metabolic and cardiovascular diseases (1–5). Since the Industrial Revolution, modern advancements in agriculture, transportation, and manufacture have had a profound impact on human diets and activity patterns, such that calorie-dense food is often easily accessible and adequate nutrition can be achieved with a sedentary lifestyle. This state of affairs, which is typical in western societies but spreading across developing countries, stands in stark contrast to the ecological conditions humans experienced through most of our species’ evolutionary history. Consequently, the ‘mismatch’ between human physiology – which evolved to cope with a mixed plant- and meat-based diet, activity-intensive foraging, and periods of resource scarcity – and western, industrialized lifestyles has been hypothesized to explain the current epidemic of cardio-metabolic disease (6–9).

In an effort to understand cardio-metabolic disease from an evolutionary perspective, researchers have studied small-scale, subsistence-level societies (e.g., hunter-gatherers, forager-horticulturalists, and pastoralists) whose diets and activity patterns are more in line with their evolutionary history (6). In particular, modern-day hunter-gatherers practice a lifestyle that is arguably representative of the distant human evolutionary past (∼300k years) (10), while forager-horticulturalists and pastoralists rely on plant and animal domestication practices that evolved ∼12k years ago (11). These populations are thus ‘matched’ to their evolutionary past on different time scales, yet all studied subsistence-level populations show extremely low levels of type II diabetes, hypertension, obesity, and heart disease relative to the US and Europe (12–18). In further support of an association between evolutionary mismatch/industrialization and cardio-metabolic disease, small-scale, indigenous populations transitioning toward market-based economies report much higher rates of obesity and metabolic syndrome than populations that have remained subsistence-level (19, 20). Similarly, individuals living in isolated, rural areas in developing countries exhibit lower levels of hypertension, type II diabetes, and obesity, relative to urban individuals with access to modern amenities and the market economy (15, 21–24).

Taken together, current evidence thus shows that as populations transition away from subsistence-level lifestyles, cardio-metabolic health declines. These patterns superficially support the evolutionary mismatch hypothesis, but cannot be disentangled from the idea that environmental, dietary, or lifestyle factors that differ between subsistence-level and urban, industrialized populations impact cardio-metabolic health regardless of a population’s evolutionary history. In other words, it is unclear whether hunter-gatherers, horticulturalists, and pastoralists are healthier than their urban, industrialized counterparts because of the ‘match’ between their lifestyle and evolved physiology, or more simply because some aspect of these lifestyles are beneficial and would be for any individual regardless of their evolved physiology. Because these ideas cannot be disentangled, we discuss evolutionary mismatch, industrialization, urbanization, and market-integration (i.e., an increased reliance on the market economy) as parallel explanations for cardio-metabolic disease.

Despite the lessons learned from studying subsistence-level populations thus far, three major gaps remain and limit our understanding of how industrialization increases disease risk. First, no large-scale health comparisons between small-scale and industrialized societies, or between urban and rural residents within a given country, have contrasted individuals from a single genetic background, making it difficult to disentangle genetic versus environmental contributions to health. Second, while chronic disease risk seems to track *degrees* of mismatch and industrialization (e.g., rates of type II diabetes follow a rank order of US, urban areas in developing countries, and small-scale societies (6, 23–25)), most studies have only compared two populations or lifestyle groups with many confounded environmental differences. Thus, it has been difficult to identify the shape of the industrialization-health relationship, or the specific lifestyle factors responsible for increasing cardio-metabolic disease risk. Third, research to date has focused almost exclusively on the consequences of exposure to industrialized environments in adulthood (26), despite mounting evidence that early life conditions can profoundly impact long-term health (27, 28). Because these influences remain unaccounted for, it is unclear how lifestyle effects across the life course interact or accumulate to determine adult health.

To address these gaps, we collected interviews and cardio-metabolic health biomarker data from the Turkana – a subsistence-level, pastoralist population from a remote desert in northwest Kenya (29, 30) (Figure 1). The Turkana and their ancestors have practiced nomadic pastoralism in arid regions of East Africa for thousands of years (30), and present-day, traditional Turkana still rely on livestock for subsistence: 70-80% of calories are derived from animal products (29). However, as infrastructure in Kenya has improved in the last few decades, small-scale markets have expanded into northwest Kenya leading some Turkana to no longer practice nomadic pastoralism; instead, they make and sell charcoal or woven baskets, or keep animals in a fixed location for trade rather than subsistence. Further, in addition to the emergence of this ‘non-pastoralist’ (but still relatively subsistence-level) subgroup, some individuals have left the Turkana homelands entirely and now live in highly urbanized parts of central Kenya (Figure 1). The Turkana situation thus presents a unique opportunity, in that individuals of the same genetic background can be found across a substantial lifestyle gradient ranging from more ‘matched’ to ‘mismatched’ with their traditional practices. Further, because many Turkana are currently migrating between rural and urban areas within their lifetime, we are able to test how environmental variation at each life stage independently contributes to adult health outcomes.

**Figure 1.**
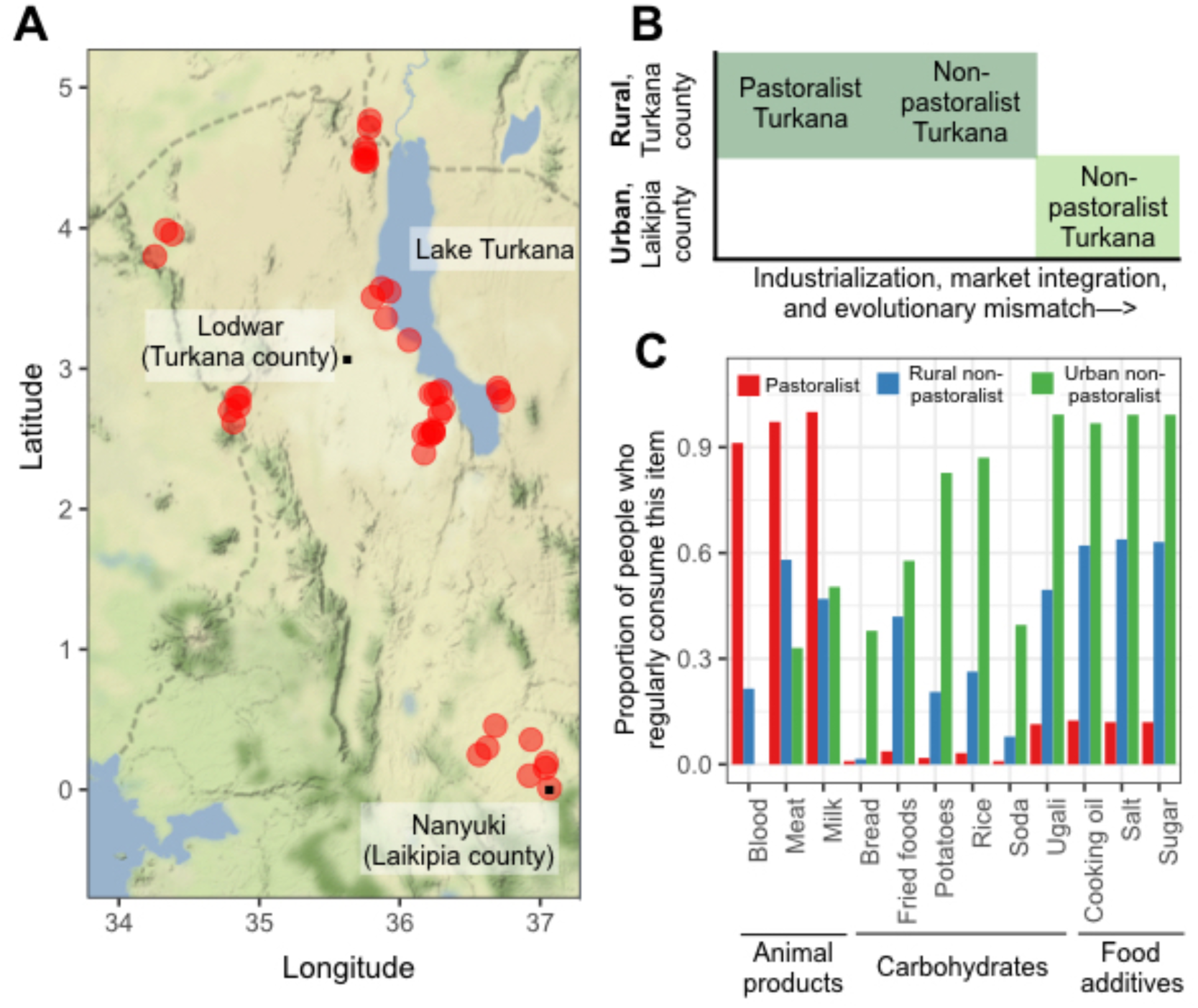
Sampling and dataset overview. (A) Sampling locations throughout northern and central Kenya are marked with red dots, the county borders are marked with dashed lines. In both Laikipia and Turkana counties, the largest city (which is generally central within each county) is marked with a black dot. (B) Schematic describing the three lifestyle groups that were sampled as part of this study. (C) The proportion of people from each lifestyle group who reported that they consumed a particular item ‘regularly’, defined as ‘1-2 times per week’, ‘>2 times per week’, or ‘every day’. People who reported that they consumed a particular item ‘rarely’ or ‘never’ were categorized as not consuming the item regularly. Animal products are a staple of the traditional pastoralist diet (66), while carbohydrates and food additives – which can only be obtained through trade – are indicative of market-integration.

Capitalizing on this unique opportunity, we sampled 1226 adult Turkana in 44 locations from the following lifestyle groups: (i) individuals practicing subsistence-level pastoralism in the Turkana homelands, (ii) individuals that do not practice pastoralism but live in the same remote, rural area, and (iii) individuals living in urban locations in central Kenya (Figure 1). We found that cardio-metabolic profiles across 10 biomarkers were favorable in pastoralist Turkana, and rates of obesity and metabolic syndrome were low, similar to other subsistence-level populations (12–18). Comparisons within the Turkana revealed a non-linear relationship between the extent of industrialization or evolutionary mismatch and cardio-metabolic health: no significant biomarker differences were found between pastoralists and non-pastoralists from rural areas. However, we found strong, sometimes sex-specific, differences in health between these two groups and non-pastoralists living in urban areas, though metabolic dysfunction among urban Turkana did not reach levels observed in the US. Using formal mediation analyses (31, 32), we show that carbohydrate consumption and indices of market integration may explain health shifts in urban Turkana. Finally, we show that a proxy of urbanization (population density) experienced around the time of birth is associated with worse adult metabolic health, independent of adult lifestyle. In other words, the health consequences of early life and adult conditions are additive and will thus accumulate across the life course.

## RESULTS

### Traditional, pastoralist Turkana are at low risk for cardio-metabolic disease

To characterize the health of the Turkana people, we collected extensive interview and biomarker data from adult Turkana sampled throughout Kenya (Figure 1). We measured body mass index (BMI), waist circumference, total cholesterol, triglycerides, high and low density lipoproteins (HDL, LDL), body fat percentage, systolic and diastolic blood pressure, and blood glucose levels (see Table S1 for biomarker-specific sample sizes). We also created a composite measure of health, defined as the proportion of measured biomarkers for a given individual that exceed cutoffs set by the CDC or the American Heart Association as being indicative of disease (see SI Materials and Methods).

As has been observed in other subsistence-level populations, we found extremely low levels of cardio-metabolic disease among traditional, pastoralist Turkana: no individuals met the criteria for obesity (BMI>30) or metabolic syndrome (33), and only 6.4% of individuals had hypertension (blood pressure > 135/85 (33)). Further, across 8 cardio-metabolic biomarkers that have been measured consistently in other subsistence-level populations (12–18), the mean values observed in the Turkana were generally within the range of what others have reported (Table S2 and Table S3). Mean body fat percentage (mean ± SD for females = 20.45% ± 4.57) and BMI (19.99 kg/m^2^ ± 2.14) were on the lower extremes, but were similar to other pastoralists (mean BMI in the Fulani and the Maasai = 20.2 kg/m^2^ and 20.7 kg/m^2^, respectively) and to a small study of the Turkana conducted in the 1980s (mean BMI = 17.7 kg/m^2^ (34)). Notably, the only biomarkers that were strongly differentiated in traditional, pastoralist Turkana were HDL (72.69 mg/dl ± 14.72) and LDL cholesterol levels (60.89 mg/dl ± 20.22), both of which were even more favorable than what has been observed in other subsistence-level groups (range of reported means for HDL = 34.45-49.11 mg/dl, LDL = 72.70-92.81 mg/dl).

### Pastoralist Turkana and rural non-pastoralist Turkana have similar biomarker profiles, while urban Turkana exhibit poorer metabolic health

Next, we sought to understand the shape of the relationship between industrialization/ mismatch and cardio-metabolic health within the Turkana, by comparing biomarker values across the three lifestyle categories. Using linear models controlling for age and sex, we found that Turkana practicing traditional pastoralism did not differ in any of the 10 measured biomarkers relative to non-pastoralist Turkana living in similarly rural areas (all p-values>0.05; Figure 2 and Table S5), despite there being major dietary difference between these groups (Figure 1 and Table S4). Pastoralist and rural non-pastoralist Turkana did significantly differ in our composite measure of metabolic health, with non-pastoralist Turkana exhibiting more biomarker values above clinical cutoffs (average proportion of biomarkers above cutoffs = 4.02% and 6.82% for pastoralists and non-pastoralists, respectively, p-value=1.39×10^−3^, FDR<5%; Figure 2).

**Figure 2.**
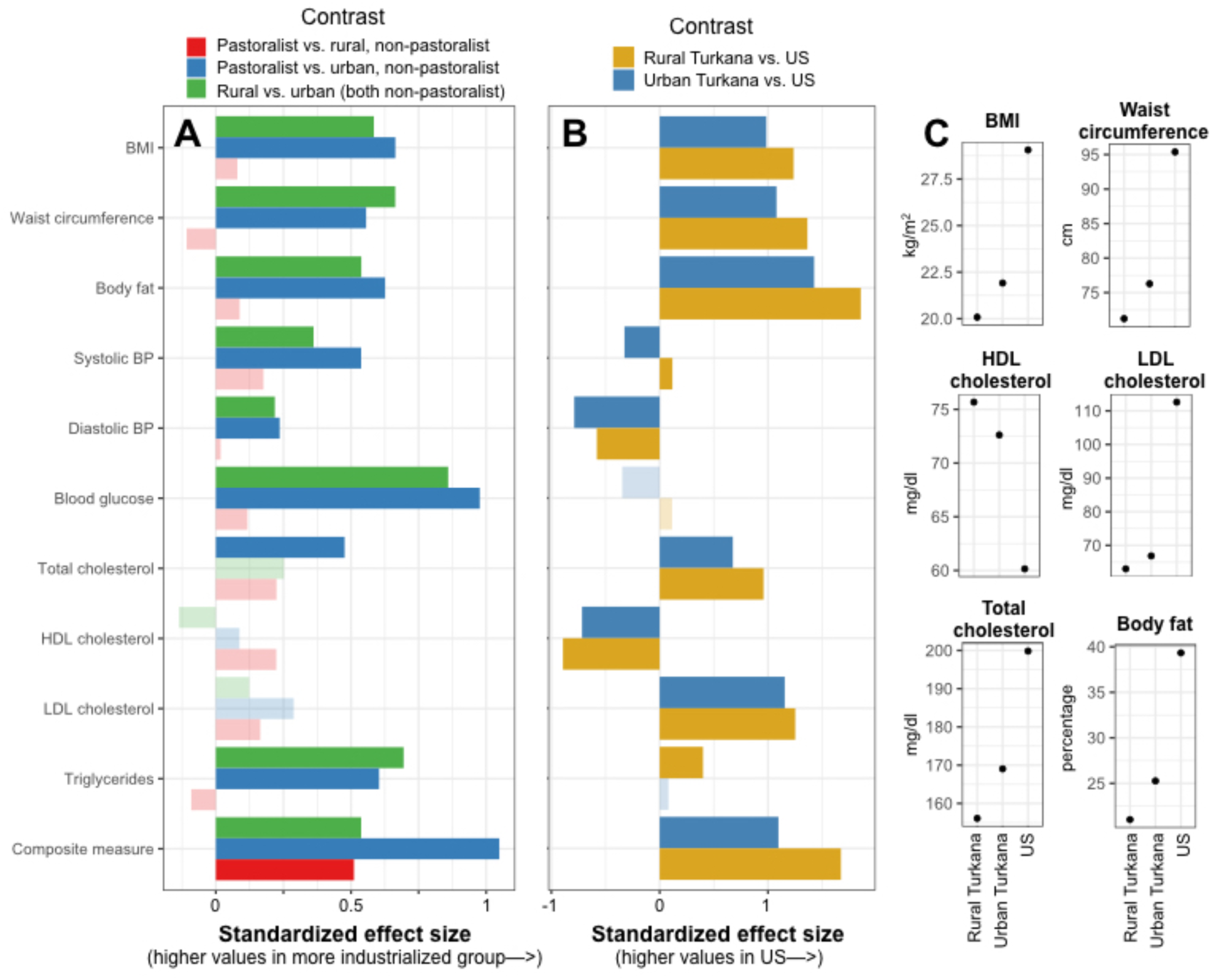
Pastoralist and rural non-pastoralist Turkana have similar health profiles, while biomarkers of metabolic dysfunction are elevated in urban Turkana but not as extremely as in the US. (A) Effect sizes for contrasts between pastoralist, rural non-pastoralist, and urban non-pastoralist Turkana (from linear models controlling for age and sex; Table S5). Effect sizes are standardized, such that the x-axis represents the difference in terms of standard deviations between groups. (B) Standardized effect sizes for contrasts between rural Turkana (pastoralist and rural non-pastoralist grouped together), urban non-pastoralist Turkana, and the US (from linear models controlling for age and sex; Table S6). In panels A and B, transparent bars represent effect sizes that were not significant (FDR>5%), and analyses of body fat and blood glucose focus on females only (see SI Materials and Methods). (C) Predicted values for a typical rural Turkana, urban Turkana, and US individual are shown for a subset of significant biomarkers. Estimates were obtained using coefficients from fitted models, for a female of average age (see SI Materials and Methods).

Strikingly, biomarker values for both pastoralist Turkana and non-pastoralist, rural Turkana were consistently more favorable than those observed in non-pastoralist Turkana living in urban areas in central Kenya. People living in urban areas exhibited composite measures indicative of worse cumulative health (average proportion of biomarkers above cutoffs = 13.42%), higher BMIs and body fat percentages, larger waist circumferences, higher blood pressure, and higher levels of total cholesterol, triglycerides, and blood glucose (all FDR<5%; Figure 2 and Table S5). In fact, the only tested variables that did not exhibit differences between urban and rural Turkana (both pastoralists and non-pastoralist) were HDL and LDL cholesterol levels, which were favorable in all Turkana regardless of lifestyle (Table S2 and Table S5). Using standardized effect sizes, we found that the biomarkers that differed most between the two rural groups and urban residents were blood glucose, triglycerides, and BMI (Figure 2). For example, the average urban Turkana resident has a 9.69% and 8.43% higher BMI relative to pastoralist and non-pastoralist rural Turkana, respectively.

For all 11 measures, we explored the possibility of a sex x lifestyle category interaction to determine whether men and women are differentially affected by lifestyle changes, but found that inclusion of this term only improved model fit for blood glucose levels (p-value from a likelihood ratio test=1.838×10^−4^) and body fat percentage (p-value=4.234×10^−3^; Table S5). In both cases, women experienced lifestyle effects on health, while men did not (Figure S1). Importantly, these results agree with several previous studies, which have shown that females are often more sensitive than males to lifestyle effects on cardio-metabolic health biomarkers (4, 16, 20, 23). Whether this increased sensitivity is due to unique aspects of female biology, or to consistent, sex-targeted changes in lifestyle that occur when populations become more industrialized, remains unclear and an area ripe for future study.

### Biomarker profiles are more favorable among both rural and urban Turkana relative to the US

We next asked whether the biomarker levels observed among urban Turkana approached those observed in a fully western, industrialized society (specifically, the US). We note that a caveat of these analyses is that they must include different genetic backgrounds, since Turkana individuals are not found in fully industrialized countries.

To compare metabolic health between the US, rural Turkana (grouping pastoralists and non-pastoralists, since these groups were minimally differentiated in previous analyses), and urban Turkana, we combined biomarker data from the CDC’s National Health and Nutrition Examination Survey (NHANES) conducted in 2006 (35) with our Turkana data, and used linear models to test for biomarker differences. These comparisons revealed that, while urban Turkana exhibit biomarker values indicative of poorer health than rural Turkana, urban Turkana have more favorable metabolic profiles than the US (Figure 2, Figure S2-S3, and Table S6). This pattern held for all measures except (i) blood glucose levels, where no differences were observed (p-value for US versus rural Turkana=0.166, US versus urban Turkana=0.074); (ii) triglycerides, where urban Turkana could not be distinguished from the US (p-value=0.627); (iii) systolic blood pressure, where urban Turkana exhibited higher values than the US (4.55% higher, p=3.02×10^−6^, FDR<5%), and (iv) diastolic blood pressure, where mean values for both urban and rural Turkana were surprisingly higher than the US (US versus rural Turkana: 8.77% higher, p=4.43×10^−65^; US versus urban Turkana: 11.69% higher, p=3.58×10^−31^, FDR<5% for both comparisons; Figure S4). These differences in diastolic blood pressure remained after removing all US individuals taking cardio-metabolic medications (US versus rural Turkana: 6.91% higher, p=3.27×10^−67^; US versus urban Turkana: 10.37% higher, p=2.21×10^−35^). However, two pieces of evidence suggest the higher diastolic blood pressure values observed in the Turkana are not pathological: (i) values for rural Turkana (77.43 mm Hg ± 15.22) are similar to estimates from other subsistence-level populations (range of published means = 70.9-79.9 mm Hg; Table S2) and (ii) far fewer rural Turkana meet the criteria for hypertension relative to the US (Figure S3).

For measures that exhibited differences between urban Turkana and the US in the expected directions, these effect sizes were consistently much larger in magnitude than the differences we observed between rural and urban Turkana (Figure 2). For example, while the average urban Turkana experiences a 9% increase in BMI relative to their rural counterparts, the average US individual has a BMI that is 44% and 32% higher than rural and urban Turkana, respectively. Similarly, while the average proportion of biomarkers above clinical cutoffs is 6.22% in rural Turkana and 13.42% in urban Turkana, this number rises to 38.84% in the US.

### Health shifts in urban Turkana are weakly mediated by increased consumption of market carbohydrates, and more strongly by indices of urbanicity

A major challenge in health research is to identify the specific dietary, lifestyle or environmental inputs that explain differential outcomes between urban and rural or industrialized and non-industrialized populations (16, 20). To do so, we turned to interview data collected for each individual (see SI Materials and Methods), which revealed substantial differences in diet, urbanicity, and market access in rural versus urban Turkana (Figure 1 and Figure 3). We paired these interview data with mediation analyses originally developed in the social sciences (31, 32), to formally test whether the effect of a predictor (X) on an outcome (Y) is direct, or is instead indirectly explained by a third variable (M) such that X→M→Y (Figure 3). Using this statistical framework, we tested whether increased consumption of market-based carbohydrates (e.g., soda, bread, rice), reduced consumption of animal products (e.g., blood, milk, meat), poorer health habits, ownership of more market-based goods and modern amenities (e.g., cell phone, finished floor, electricity), occupation that is more market-integrated, and residence in a more populated or developed area (measured via population density, distance to a major city, and female education levels) could explain the decline in metabolic health observed in urban Turkana (see SI Materials and Methods). In particular, we predicted that lifestyle effects on health would be mediated by a shift toward a diet that incorporates more carbohydrates and fewer animal products in urban Turkana. These analyses focused on biomarkers for which our sample sizes were the largest, since dietary data was not available for all individuals (see Table S7 for sample sizes).

**Figure 3.**
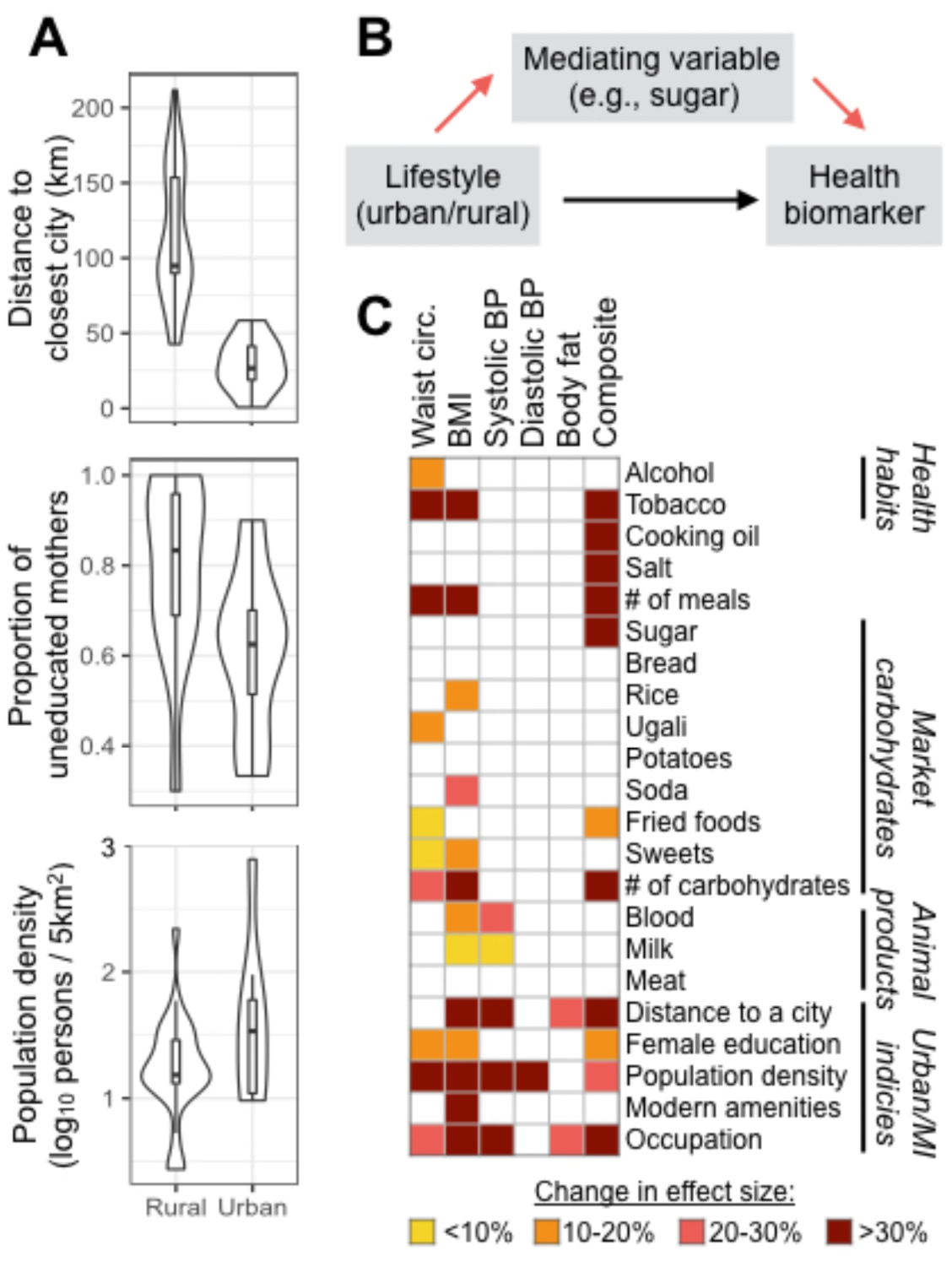
Urban-rural health differences are only weakly mediated by diet. (A) Key measures of urbanicity and market-integration used in mediation analyses, with means and distributions shown for urban and rural Turkana. (B) Schematic of mediation analyses. Specifically, mediation analyses test the hypothesis that lifestyle effects on health are explained by an intermediate variable, such as consumption of particular food items (red arrows); alternatively, lifestyle effects on health may be direct (black arrow) or mediated by a variable we did not measure. (C) Summary of mediation analysis results, where colored squares indicate a variable that was found to significantly explain urban-rural health differences in a given biomarker. Significant mediators are colored based on how much the lifestyle effect (urban/rural) decreased when a given mediator was included in the model. MI = market-integration. Full results and samples sizes for mediation analyses are presented in Table S7.

In support of our predictions, urban-rural differences in waist circumference, BMI, and our composite measure of health were mediated by increased consumption of mostly refined, market-derived carbohydrates (e.g., sugar, soda, fried foods) and decreased consumption of animal products (milk and blood) in urban Turkana (Figure 3 and Table S7). Notably, a tally of the number of different carbohydrate items a given individual consumed was a strong and consistent predictor across these three biomarkers, suggesting that individual dietary components may matter less than overall exposure to refined carbohydrates. Contrary to our predictions, we did not find that dietary differences mediated urban-rural differences in systolic blood pressure, diastolic blood pressure, or body fat percentage. Instead, these measures were explained by variables that captured how industrialized and market-integrated a given individual’s lifestyle was, which was also important for waist circumference, BMI, and our composite measure of health in addition to dietary effects. For example, fine-scale measures of population density significantly mediated 5/6 tested biomarkers, as did variation in how reliant on the market economy an individual’s occupation was (Figure 3 and Table S7). Further, these indices of urbanicity and market-integration tended to be stronger mediators than dietary variables (Figure 3).

To understand the degree to which the mediators we identified explain the relationship between lifestyle and a given biomarker, we compared the magnitude of the lifestyle effect in our original models (controlling for age and sex, without any mediators) to the effect estimated in the presence of all significant mediators. If the mediators fully explain the relationship between lifestyle and a given biomarker, we would expect the estimate of the lifestyle effect to be 0 in the second model. These analyses revealed that the mediators we identified explain most of the relationship between lifestyle and waist circumference (effect size decrease = 90.7%), BMI (79.9%), systolic blood pressure (74.9%), and composite health (64.1%), but explain only a small portion of lifestyle effects on diastolic blood pressure (10.0%) and body fat (23.5%; Table S7).

### Cumulative exposure to urban environments across the life course compromises metabolic health

Finally, we were interested in understanding whether being born in an urban, industrialized environment had long-term effects on health, above and beyond the effects of adult lifestyle we had already identified. We were motivated to look for early life effects because work in humans and non-human animals has demonstrated strong associations between diet or ecology during the first years of life and fitness-related traits measured many years later (28, 36–38). Two major hypotheses have been proposed to explain why this ‘embedding’ of early life conditions into long-term health occurs. First, the ‘predictive adaptive response’ (PAR) hypothesis posits that organisms adjust their phenotype during development in anticipation of predicted adult conditions. Individuals that encounter adult environments that ‘match’ their early conditions are predicted to gain a selective advantage, whereas animals that encounter ‘mismatched’ adult environments should suffer a fitness cost (27, 38–41). In contrast, the ‘developmental constraints’ (DC) or ‘silver spoon’ hypothesis predicts a simple relationship between early environmental quality and adult fitness: individuals born in high-quality environments experience a fitness advantage regardless of the adult environment (38, 42, 43). Importantly, under DC, poor-quality early life conditions cannot be ameliorated by matching adult and early life environments, instead, the effects of environmental adversity accumulate across the life course.

We found no evidence that individuals who experienced matched early life and adult environments had better metabolic health in adulthood than individuals who experienced mismatched early life and adult conditions (p>0.05 for all biomarkers). In particular, we tested for interaction effects between the population density of each individual’s birth location (estimated for their year of birth) and a binary factor indicating whether the adult environment was urban or rural (Table S8; see Table S9 for parallel analyses using population density to define the adult environment as a continuous measure). This analysis was possible given the within-lifetime migrations of many Turkana between urban and rural areas: only 19.52% and 33.01% of urban and rural Turkana, respectively, were sampled within 10km of their birthplace, and the correlation between birth and sampling location population densities was low (R^2^ = 0.115, p<10^−16^; Figure S5).

While we observed no evidence for interaction effects supporting PAR, we did find strong main effects of early life population density on adult waist circumference (p=7.33×10^−8^), BMI (p=1.35×10^−7^), body fat (p=1.57×10^−5^), diastolic blood pressure (p=2.01×10^−2^), and our composite measure of health (p=3.40×10^−4^; all FDR<5%), in support of DC. For all biomarkers, being born in a location with high population density was associated with poorer adult metabolic health (Figure 4 and Table S8), and the early life environment effect was on the same order of magnitude as the effect of living in an urban versus rural location in adulthood (see Table S8 and SI Materials and Methods). For example, BMIs are 5.69% higher in urban versus rural areas, while individuals born in areas from the 25^th^ versus 75^th^ percentile of the early life population density distribution exhibit BMIs that differ by 3.34%. Similarly, the effect of the adult environment (urban compared to rural) on female body fat percentages is 11.14%, while the effect of early life population density (25^th^ compared to 75^th^ percentile) is 20.7% (Table S8).

**Figure 4.**
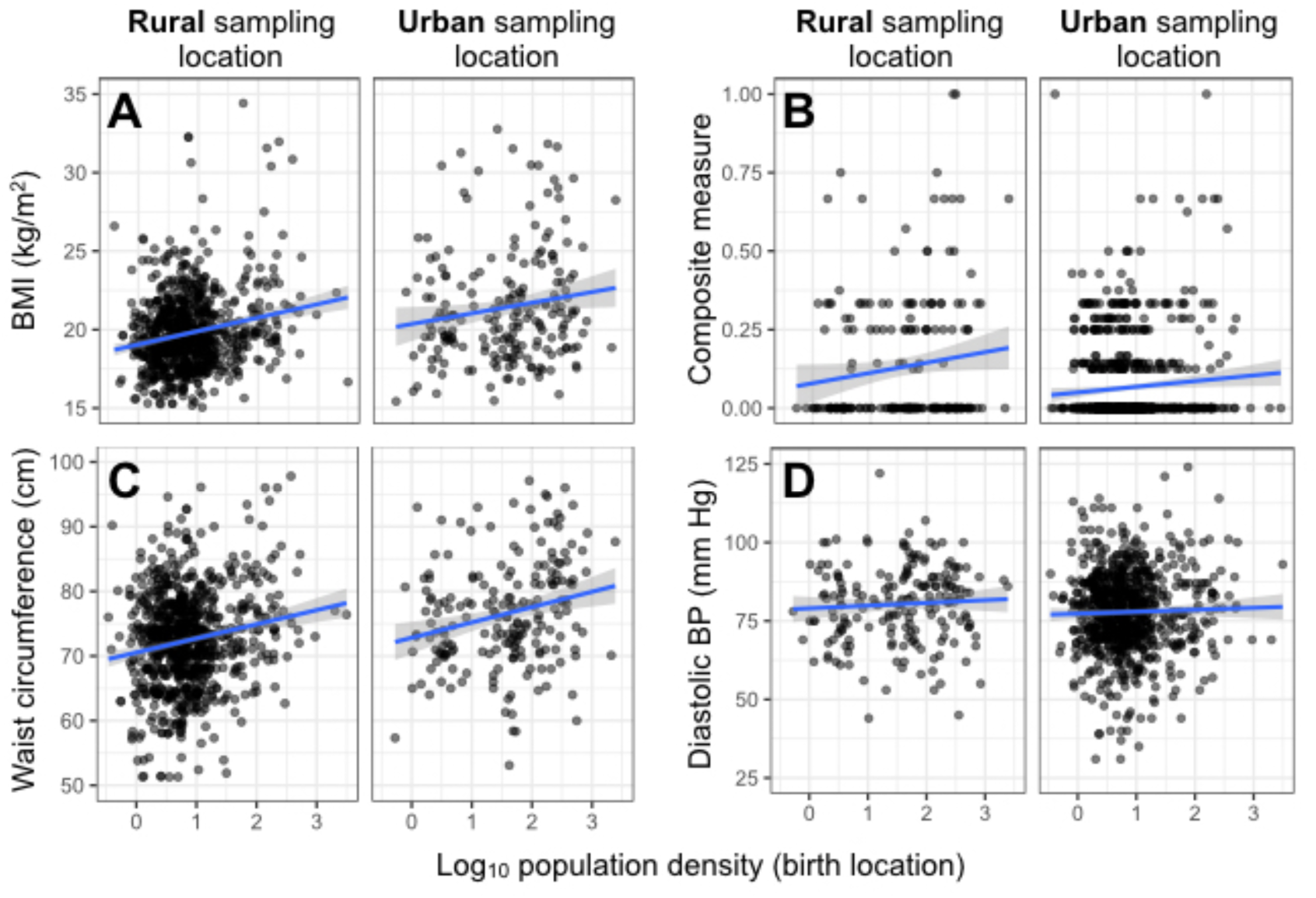
Early life population density predicts biomarkers of adult health. The relationship between population density of each individual’s birth location and (A) BMI, (B) our composite measure of health, (C) waist circumference, and (D) diastolic blood pressure are shown for individuals sampled in rural and urban locations, respectively. Notably, while the intercept for a linear fit between early life population density and each biomarker differs between rural and urban sampling locations (indicating mean differences in biomarker values as a function of adult lifestyle), the slope of the line does not. In other words, we find no evidence that the relationship between early life conditions and adult is contingent on the adult environmnt (as predicted by PAR). Instead, being born in an urban location predicts poorer metabolic health regardless of the adult environment.

## DISCUSSION

By sampling a single genetic background across a substantial lifestyle gradient, we show that (i) traditional, pastoralist Turkana exhibit low levels of cardio-metabolic disease and (ii) increasing industrialization – in both early life and adulthood – has detrimental effects on metabolic health. While our study cannot definitively address whether compromised health in urban Turkana is driven by industrialized environments, or more specifically by mismatches between the Turkana’s evolved traits and these conditions, we provide more direct evidence for the latter hypothesis than existing work that confounds lifestyle and genetic background. Truly separating evolutionary mismatch from main effects of industrialization would require, for example, placing individuals with a different evolutionary history into our studied lifestyle groups. Under mismatch, we would expect Turkana to exhibit better health than individuals that are not locally adapted to nomadic, desert pastoralism when both groups engage in this lifestyle. Under a hypothesis of main environmental effects, we’d expect similar health differences between individuals practicing pastoralism versus living in urban areas, regardless of genetic background and evolutionary history. However, without the ability to perform transplant experiments in humans, studying a lifestyle gradient ranging from well-matched to mismatched with the known evolutionary history of a population – as we have done in the Turkana – is arguably the best natural test of the evolutionary mismatch hypothesis.

Our observation that pastoralist Turkana do not suffer from cardio-metabolic diseases common in western societies agrees with work in subsistence-level hunter-gatherers, forager horticulturalists, and pastoralists (12–18). More generally, it supports previous conclusions that many types of mixed plant- and meat-based diets are compatible with cardio-metabolic health (6), and that mismatches between the distant human hunter-gatherer past and the subsistence-level practices of forager horticulturalists or pastoralists do not lead to disease (44). In other words, contemporary hunter-gatherers are most aligned with human subsistence practices that evolved ∼300k years ago (10), but they do not exhibit better cardio-metabolic health relative to forager horticulturalists or pastoralists, whose subsistence practices evolved ∼12k years ago (11) (Table S2-S3). Instead, we find evidence consistent with the idea that extreme mismatches between the recent evolutionary history of a population and lifestyle are needed to produce health declines; in the Turkana, this situation manifests in urban, industrialized areas but not in rural areas with limited access to the market economy.

Our study thus joins other work that has analyzed the health consequences of industrialization across multiple genetic backgrounds (12–18, 21, 22, 45), or more limited lifestyle gradients (16, 20, 46–48), in concluding that the environmental conditions typical of western societies put individuals at risk for cardio-metabolic disease. Importantly, because our study assessed health in individuals who experience no, limited, or substantial access to the market economy, we were able to determine that industrialization has non-linear effects on health. In particular, we find no differences between pastoralists and non-pastoralist in rural areas for 10/11 variables (Figure 2), despite non-pastoralists consuming processed carbohydrates that are atypical of traditional practices (Figure 1 and Table S2). Nevertheless, rural non-pastoralists still live in remote areas, engage in activity-intensive subsistence activities, and rely far less heavily on markets than urban Turkana. Our results suggest that this type of lifestyle, while different from traditional pastoralism, has not crossed the threshold necessary to produce cardio-metabolic health issues. More generally, this ‘threshold model’ may help explain heterogeneity in previous studies, where small degrees of mismatch and market-integration have produced inconsistent changes in cardio-metabolic health biomarkers (16, 49, 50).

While our dataset does not capture all of the variables that mediate urban-rural health differences in the Turkana, we were able to account for a substantial portion (>60%) of lifestyle effects on waist circumference, BMI, systolic blood pressure, and composite health. In line with our expectations, increases in these biomarkers in urban areas was mediated by greater reliance on refined carbohydrates and reduced consumption of animal products (Figure 1 and Figure 3). However, our mediation analyses also show that measures of how industrialized and market-integrated an individual’s lifestyle is (e.g., population density, distance to a major city, and female education levels) have stronger explanatory power across a greater number of biomarkers than diet. It is likely that these indices serve as proxies for unmeasured, more proximate mediators – such as psychosocial stress, nutrient balance, total caloric intake, or total energy expenditure from physical activity – all of which vary as a function of industrialization and can affect components of health (6, 18, 51–54). The fact that the number of meals eaten per day (which is typically 1 in rural areas and 2-3 in urban areas) was a strong mediator for 3/6 variables points toward a potential role for total caloric intake, while the importance of occupation suggests activity levels are also probably key. Ongoing work with the Turkana will address these unmeasured sources of variance.

In addition to the pervasive influence of adult lifestyle on metabolic physiology we observe in the Turkana, our analyses also revealed appreciable effects of early life environments.

In particular, controlling for the adult environment (urban or rural), birth location population density was a significant predictor of BMI, waist circumference, diastolic blood pressure, our composite measure of health, and body fat. Further, the impact of early life and adult conditions appear to be on the same order of magnitude: 2-6% differences in BMI, waist circumference, and diastolic blood pressure are observed as a function of each life stage, while body fat and our composite measure show changes in the 11-20% range (Table S8).

Importantly, we did not find evidence that individuals who grew up in rural versus urban conditions were more ‘prepared’ for these environments later in life, as predicted by PAR. These findings agree with work in pre-industrial human populations and long-lived mammals, which have found weak or no support for PARs (55–59). This is potentially because early life ecological conditions are often a poor predictor of adult environments for long-lived organisms; consequently, ‘matching’ individual physiology to an unpredictable adult environment is a poor strategy that is unlikely to evolve (60–62). Instead, our work joins that of others in concluding that challenging early life environments simply incur long-term health costs (55–59). In the Turkana, greater overall exposure to industrialized environments across the life course is associated with the largest health burdens; understanding the generality of these effects will require longitudinal, life course data from other populations undergoing industrial transitions (e.g., (26)). Further, additional effort should be spent characterizing early life environments and potential mediating variables with the same attention that has been paid to adult conditions.

## METHODS

### Sampling overview

Data were collected between April 2018 and March 2019 in Turkana and Laikipia counties in Kenya. During this time, researchers visited locations where individuals of Turkana ancestry were known to reside (Figure 1). At each sampling location, healthy adults (>18 years old) of self-reported Turkana ancestry were invited to participate in the study, which involved a structured interview and measurement of 10 cardio-metabolic biomarkers. Participation rates of eligible adults were high (>75%). GPS coordinates were recorded on a handheld Garmin GPSMAP 64 device at each sampling location.

This study was approved by Princeton University’s Institutional Review Board for Human Subjects Research (IRB# 10237), and Maseno University’s Ethics Review Committee (MSU/DRPI/MUERC/00519/18). We also received county-level approval from both Laikipia and Turkana counties for research activities, as well as research permits from Kenya’s National Commission for Science, Technology and Innovation (NACOSTI/P/18/46195/24671). Written, informed consent was obtained from all participants, after the study goals, sampling procedures, and potential risks were explained to them in their native language (by both a local official, usually the village chief, and by researchers or field assistants).

### Statistical analyses

For each of the 11 measures (10 direct measures and 1 derived, composite measure; Table S1), we used linear models controlling for age and sex to test for lifestyle effects on health within the Turkana (comparing pastoralists, rural non-pastoralists, and urban non-pastoralists). We also used a likelihood ratio test to determine whether a sex x lifestyle interaction effect improved the fit of each model; these analyses revealed that body fat percentage and blood glucose levels exhibited lifestyle effects in females only, so downstream analyses of these two biomarkers excluded males. To understand whether cardio-metabolic profiles in urban Turkana were as extreme as what is observed in the US, we downloaded data from the CDC’s National Health and Nutrition Examination Survey (NHANES) conducted in 2006 (35) and filtered for adults aged 18-65. We used these data as input in linear models, controlling for age and sex, to test for differences between rural Turkana (grouping pastoralist and non-pastoralist), urban Turkana, and the US. In all analyses, we corrected for multiple hypothesis testing using a Benjamini–Hochberg false discovery rate (FDR) (63). We considered a given lifestyle contrast to be significant if the FDR-corrected p-value was less than 0.05 (equivalent to a 5% FDR threshold).

To identify dietary and lifestyle factors that explain biomarker variation between urban and rural Turkana, we conducted formal mediation analyses (31, 32). Specifically, we compared the estimate of the urban/rural effect on waist circumference, BMI, diastolic and systolic blood pressure, body fat, and composite health estimated from linear models that included versus excluded each potential mediator (all potential mediators are listed in Table S7). For each biomarker-mediator pair, we used 1000 iterations of bootstrap resampling to assess significance, and considered a variable to be a significant mediator if the lower bound of the 95% confidence interval (for the change in effect size between the model with and without the mediator) did not overlap with 0.

To understand whether early life population density affected adult health, we modeled each biomarker as a function of age, sex, population density during the year of birth (estimated using data from NASA’s Socioeconomic Data and Applications Center (SEDAC; https://doi.org/10.7927/H49C6VHW), a binary variable describing the adult environment (urban/rural), and the interaction between early life and adult conditions. This modeling strategy is similar to the approach of (55–58, 64), and has been previously used to test whether the health consequences of early life conditions are contingent on the adult environment (as predicted by PAR) or are instead additive and independent (as predicted by DC). In cases where we found no evidence for significant interactions between early life and adult conditions (p>0.05), we removed this term and report effect sizes and p-values from linear models with only independent effects of early life and adult environments.

All statistical analyses were performed in R (65). Further details for all sampling and statistical procedures can be found in the SI Materials and Methods.

## ACKNOWLEDGMENTS

This work was supported by an award to JFA through Princeton University’s Dean for Research Innovations Funds. AJL was supported by a postdoctoral fellowship from the Helen Hay Whitney Foundation. We thank Simon Lowasa, Charles Waigwa, Francis Lotukoi, and Erick Loowoth for their invaluable assistance with data collection, as well as Denis Misiko Mukhongo, Susan Ngatia, and Benjamin Mbau for their contributions to the project. We thank Michelle Ndegwa for expert database and project management, and Jeremy Orina and Dan Rubenstein for logistical help in Kenya. We are also grateful to the staff of Mpala Research Centre for their essential support, especially Fardosa Hassan, Cosmas Nzomo, Beatrice Wanjohi, Gikenye Chege, Tony Maina, and Julius Nakolonyo. Finally, thanks to the Ayroles and Graham labs, as well as Jeanne Altmann, for feedback on this project.

## DATA AVAILABILITY

Data underlying the analyses presented in this paper will be deposited on Dryad and made available after publication.

## AUTHOR CONTRIBUTIONS

AJL, DM, JFA designed research; AJL, DM, JK, JFA performed research; AJL analyzed data; and AJL, MG, JFA wrote the paper, with contributions from all co-authors.

