## Supplementary Methods and Figures for "Urbanization and market-integration have strong, non-linear effects on cardio-metabolic health in the Turkana"

Table of contents:

1. Supplementary Materials and Methods

2. Supplementary Figures

- Figure S1. Mean biomarker values for measures with a significant sex x lifestyle interaction.
- Figure S2. Mean biomarker values for Turkana individuals versus the US (NHANES).
- Figure S3. Proportion of Turkana versus US (NHANES) individuals whose biomarker values exceed clinical cutoffs.
- Figure S4. Blood pressure values for Turkana individuals versus the US (NHANES).
- Figure S5. Relationship between birth location and location at time of sampling.
- Figure S6. Correlation coefficients among measured biomarkers.

3. Supplementary References

### Supplementary Materials and Methods

#### Study population

The Turkana are thought to be descendants of the Jie people of Uganda based on their linguistic and cultural similarity. The ancestors of the present day Turkana likely entered Kenya in the early 18th century, and expanded to their current range across Turkana county by ~1900 (1). Today, the Turkana represent the second largest pastoralist group in Kenya after the Maasai. Turkana county is a semi-arid desert characterized by low annual rainfall (~200 mm per year), frequent droughts, and high year round temperatures (mean day time high = 97F) (2). Most of the annual precipitation occurs only during the rainy season (March-May).

The Turkana are traditionally nomadic pastoralists; they herd five species of livestock (dromedary camels, zebu cattle, fat tailed sheep, goats, and donkeys) and rely on these animals as their main mode of subsistence (3). As a result of their pastoralist lifestyle, the Turkana have a remarkably protein-rich diet: 62% of calories are derived from milk or milk products, and 70-80% of calories are derived from animal products of some sort (3). Daily protein intake exceeds the FAO/WHO requirements by >300%, despite total caloric intake being limited (1,300–1,600 kcal/day) (4). Dietary items not derived from livestock are obtained through hunting and gathering, or through trade of animal products. For detailed descriptions of the diet, climate, and lifestyle experienced by traditional, pastoralist Turkana, see work from the South Turkana Ecosystem Project (summarized in (5)).

Over the last several decades, infrastructure in Turkana county has rapidly improved, primarily due to oil discoveries in South Sudan. As a result, this previously isolated region of Kenya has become more connected with the rest of the county, allowing market goods to flow into some regions of Turkana and allowing people from Turkana to move south into more urbanized parts of Kenya. We focused on one region that has experienced an influx of Turkana people in the last few decades – Laikipia county. Laikipia is a cosmopolitan county near the equator with several large cities (Nanyuki, Nyahururu, and Rumuruti). The climate is generally cool and temperate, with rainfall throughout Laikipia county averaging from ~400-1100 mm per year depending on the specific location (6, 7). Importantly, because most of the migration between Turkana and Laikipia has occurred within the last few decades, and because it is rare for Turkana individuals to marry non-Turkana individuals, we expect very little population structure or genetic differentiation between Turkana individuals living in Turkana versus Laikipia county.

**Interview and food frequency questionnaire data**

Structured interviews were conducted with all participants, to collect basic information about demography, reproductive and health history, and lifestyle. All interviews were conducted in a language familiar to the participant (English, Turkana, or Swahili). The following select measures from the interviews are relevant to the analyses presented in the main text:

- Sex
- Self-reported age
- Where the participant was born
- Main subsistence activity, chosen from the following categories: self-employment, formal employment, petty trade, farming, animal keeping, hunting and gathering, other.
- Occupation
- Highest education level
- Number of children

- Possession of the following items: finished floor, finished room, electricity, television set, mobile phone, flush toilet, gas cooking, indoor tap water, treated water. We asked about these items to quantify how urban or modernized each individual's life was, following previous work (8–10).
- Whether the participant was currently pregnant (for women only)
- Whether the participant was currently fasting
- Number of meals eaten per day

Following previous studies (10), we also used a food frequency questionnaire to collect information about the use of a select items among the Turkana: alcohol, tobacco, meat, milk, blood, sugar, bread, salt, cooking oil, rice, ugali, potatoes, soda, fried foods, and sweets. We focused on these items because they reflect foods that are essential (meat, milk, blood) or uncommon (all carbohydrates as well as salt, and cooking oil) in the diet of pastoralist Turkana. The uncommon items can only be obtained through trade, and we therefore viewed these items as an indicator of market integration. Participants were asked how often a specific item was consumed and were given the following answer choices: never, rarely, 1-2 times per week, >2 times per week, or every day. For mediation analyses, these answers were recoded to a numeric scale from 0-4. For mediation analyses, a 'carbohydrate score' was also created by summing up the number of carbohydrate items a given participant ate regularly (regularly was defined as more often than 'never' or 'rarely'). We note that the food frequency questionnaire was designed to provide general estimates of variation in diet between individuals, rather than exact caloric or nutritional estimates. Information about the consumption of each item among Turkana practicing different lifestyles can be found in Figure 1 and Table S2.

**Biomarker measurements**

Body mass index (BMI). Weight was recorded using a portable scale, which was always placed on a hard, flat surface. Height was recorded using a Seca 213 portable stadiometer; before height measurements were recorded, participants were asked to remove any hats or hair ornaments. Weight and height were recorded to the nearest kg and cm, respectively. BMI was calculated as weight (in kg) / height (in m)$^2$.

Waist circumference. Clothing was first removed from the waist line, and the participant was asked to stand with their feet shoulder width apart and their back straight. The top of the hip bone was located by the researcher, and measuring tape was aligned with the top of the hip bone and wrapped around the participant's waist. This procedure was completed three times, and three separate measurements of waist circumference were recorded to the nearest 0.5 cm. We considered the final waist circumference value for a given participant to be the average of the three measurements.

Lipid profiles. Approximately 8mL of intravenous blood was collected into an EDTA vacutainer tube. Immediately after collection, 40ul of each sample was used to conduct a lipid panel using the CardioCheck Plus (PTS Diagnostics) following the manufacturer's instructions. Specifically, total cholesterol, HDL cholesterol, and triglycerides were measured for each participant, and LDL cholesterol was calculated by the CardioCheck Plus. Because the CardioChek Plus cannot read triglyceride values below 50 mg/dL, both triglyceride and LDL cholesterol values were not recorded for these individuals; this is because LDL cholesterol is estimated as a function of triglyceride, HDL cholesterol, and total cholesterol levels using the Friedewald Equation (LDL cholesterol = total cholesterol - HDL cholesterol – triglycerides/5).

Blood glucose levels. A drop of whole blood (~10ul) from a finger prick using a safety lancet was used to measure glucose levels. To do so, we used the OneTouch system following the manufacturer's instructions.

Body fat percentage. Body fat percentage was measured via bioelectrical impedance using the Omron HBF-306C Handheld Body Fat Loss Monitor according to the manufacturer's instructions.

Blood pressure. Arterial blood pressure was measured as systolic blood pressure and diastolic blood pressure. All participants sat in a relaxed position with their arm flat at a 90-degree angle, and with the arm relaxed and the wrist facing up. The cuff was placed around the participant's upper arm (approximately ½ inch above the elbow). Participants were asked not to speak while a trained assistant used an Omron 10 Series Wireless Upper Arm Blood Pressure Monitor to collect a single measurement.

Composite measure. We also included a composite measure of health, which tallied the number of biomarkers above clinical cutoffs for a given individual. Specifically, we counted the number of biomarkers for each participant that met the following criteria: (i) waist circumference >89 cm for women or >102 cm for men; (ii) triglyceride levels >150 mg/dL; (iii) HDL cholesterol levels <40 mg/dL for men or <50 mg/dL for women; (iv) blood pressure >135/85; (v) fasting blood glucose levels >100 mg/dL; (vi) BMI >25; (vii) LDL cholesterol levels >100 mg/dL; and (viii) total cholesterol levels > 200 mg/dL. Cutoffs for BMI, LDL cholesterol, and total cholesterol were taken from CDC recommendations. Cutoffs for the remaining biomarkers followed the criteria used by the American Heart Association to define metabolic syndrome (11).

Spearman correlation coefficients among each of the biomarkers are presented in Figure S6.

**Defining lifestyle categories**

For analyses presented in the main text, individuals were binned into the following three categories based on their sampling location, diet, and self-reported subsistence activity: (i) pastoralist Turkana were defined as individuals that reported their main subsistence activity as 'pastoralism', that drink milk every day (i.e., that rely on their livestock for subsistence), and that live in Turkana county; (ii) non-pastoralist, rural Turkana were defined as individuals that live in Turkana county but did not meet the criteria for category (i); and (iii) non-pastoralist, urban Turkana were defined as individuals that live in Laikipia county.

We arrived at these category designations based on the following observations and analyses. First, based on our own observations and on comparisons of several commonly used measures of urbanicity (8, 9) (Figure 3), we considered all locations in Turkana county to be 'rural' and all locations in Laikipia county to be 'urban' and more industrialized or market-integrated. Second, based on studies from the South Turkana Ecosystem Project (5) and our own observations, we concluded that individuals practicing traditional pastoralism are only found in Turkana county. Based on these same sources (4), we also concluded that a major marker of traditional pastoralism was regular milk consumption, suggesting that individuals truly rely on their livestock for subsistence. While some individuals classified as rural non-pastoralists reported 'animal keeping' as their main subsistence activity, these individuals generally keep animals in a fixed location and use them for trade. Additionally, these non-nomadic animal keepers do not consume milk and blood from the animals on a regular basis (34% rarely consume milk and 66% consume milk 1-2 times per week); further, 75%, 63%, and 73% of these individuals use added sugar, salt, and cooking oil regularly, which also influenced our decision to not classify them as pastoralists. To ensure that our analyses were not biased by these decisions, particularly the classification of people who practice non-nomadic animal keeping within the general group of ‘rural non-pastoralists’, we repeated the analyses in the main text after assigning these individuals to their own, fourth category. Results were quantitatively similar to what is reported in the main text and did not suggest that non-nomadic animal keepers should be considered a third group within the rural environment (Table S10).

**Data filtering and processing**

Individuals were excluded from analyses if they met any of the following criteria: (i) pregnant women; (ii) individuals with a value that was impossible for a given biomarker and deemed to be a typo; (iii) individuals whose sampling date or location did not match known sampling dates or locations, and were therefore deemed to be errors; (iv) individuals living in Turkana who did not report their main subsistence activity; (v) individuals for whom biomarker data but not interview data were recorded; and (iv) individuals without a reported gender or age.

For the early life effects analyses, we also excluded individuals that did not report a birth location, or for whom GPS coordinates for the reported birth location could not be identified on a map. Importantly, we found no evidence that individuals with missing birth locations had systematically different health profiles compared with individuals for whom a birth location could be assigned (all $p>0.05$ for linear models testing for an effect of birth location missingness on each biomarker, controlling for age and sex). Thus, though the sample sizes for our early life effects analyses are smaller than for analyses focused on current environmental/lifestyle effects (Table S8), this sample size reduction is not systematically biased in a way that is likely to affect the results.

Prior to statistical analyses, all biomarker measures (except the composite measure of health) were mean centered and scaled by their standard deviation, using the ‘scale’ function in R. Consequently, all reported effect sizes are standardized, and represent the effect of a given variable on the outcome in terms of increases in standard deviations.

**Testing for lifestyle effects on measured biomarkers**

For each of the 10 measured biomarkers (Table S1), we used the following linear model to test for effects of lifestyle controlling for covariates:

$$y_i = \beta_0 + l_i\beta_l + a_i\beta_a + s_i\beta_s + e_i \quad (1)$$

Where $y_i$ is the normalized (mean centered and scaled by the standard deviation) biomarker value for individual $i$, $l_i$ is lifestyle (pastoralist; non-pastoralist, rural; or non-pastoralist, urban), $a_i$ is age (in years), $s_i$ is sex (male or female), and $e_i$ represents residual error. To determine whether a given biomarker exhibited a lifestyle x sex interaction, we used a likelihood ratio test to compare the fit of model (1) with the following model:

$$y_i = \beta_0 + l_i\beta_l + a_i\beta_a + s_i\beta_s + (l_i * s_i)\beta_{l*s} + e_i \quad (2)$$

In model (2), $\beta_{l*s}$ represents a lifestyle x sex interaction effect. If the p-value for the likelihood ratio test comparing model (1) and model (2) was less than 0.05, we concluded that a lifestyle x sex interaction existed, and we tested for effects of lifestyle within each gender separately (controlling for age). These analyses revealed that lifestyle influences body fat levels and blood glucose levels in females but not males (Figure S1), follow up analyses for these biomarkers therefore analyzed data from females alone.

For analyses of blood glucose levels, models (1) and (2), and versions of model (1) focused on females alone, also included a covariate noting whether the individual was fasting at the time of blood collection (yes/no). For analyses of the composite measure of health, we used the same approach and the same main and interaction effects described for models (1) and (2) paired with generalized linear models with a binomial link function to accommodate count data. Specifically, the composite measure of health was modeled as the number of biomarkers that exceeded clinical cutoffs / the number of biomarkers measured for a given individual. Only individuals with 3 or more measured biomarkers were included in this analysis.

For all 11 measures (10 biomarkers and 1 composite measure), we extracted the p-values associated with the lifestyle effect ($\beta_l$) from our models, and corrected for multiple hypothesis testing using a Benjamini–Hochberg false discovery rate (12). We considered a given lifestyle contrast to be significant if the FDR-corrected p-value was less than 0.05 (equivalent to a 5% FDR threshold). The results of all final models are presented in Table S5.

**Processing and analysis of publicly available datasets**

*Biomarker measurements from other populations*

To compare biomarker levels in the Turkana to those reported for other small-scale and industrialized societies, we extracted summary data from published work for the following measures: BMI, total cholesterol, LDL cholesterol, HDL cholesterol, triglycerides, and body fat percentage. In particular, we collected biomarker information for small-scale agriculturalist and forager horticulturalist populations (Bantu, Tsimane, Shuar), other pastoralist populations or populations known to subsist on a high-protein diet (Evenki, Fulani, Masaai), and hunter-gatherers (Hazda) (10, 14–19). We also compiled information on Turkana BMI from the 1980s, from work published by the South Turkana Ecosystem Project (20). For all biomarkers, we focused on data from adults and extracted sex-specific information where possible. We extracted the mean and standard deviation of each biomarker when reported, and in some cases, calculated the standard deviation from the reported standard error value and the sample size. All comparative data is presented in Table S3, and summaries are reported in Table S2.

To understand whether metabolic profiles in urban Turkana were as extreme as what is observed in the US, we downloaded raw data from the CDC's National Health and Nutrition Examination Survey (NHANES) conducted in 2006 (13). We filtered the data to include adults from 18-65 years of age, and combined these data with our biomarker values. We then used linear models to test for effects of lifestyle (urban Turkana, rural Turkana, or US) on standardized values of each biomarker controlling for age and sex (similar to model (1)). For analyses of diastolic blood pressure, we repeated these analyses after excluding individuals on medications defined as 'cardiovascular agents' by NHANES (this includes vasodilators, ACE inhibitors, beta blockers, and diuretics commonly used to treat high blood pressure). For fasting glucose and body fat percentage, which only show lifestyle effects in female Turkana, we analyzed females alone in the combined US and Turkana dataset. For the composite measure of health, which we calculated for the NHANES data in the same way as described in *Biomarker measurements*, we used generalized linear models with a binomial link function rather than linear models. For all 11 measures, we extracted the p-value associated with the lifestyle effect, and considered a given lifestyle contrast to be significant if the FDR-corrected p-value was less than 0.05.

*Population density*

Population density estimates were downloaded from NASA's Socioeconomic Data and Applications Center (SEDAC; https://doi.org/10.7927/H49C6VHW). Specifically, we used the

Gridded Population of the World database (v4.11), which consists of estimates of human population density (number of persons per square kilometer) based on counts consistent with national censuses and population registers. These data are available for the years 2000, 2005, 2010, 2015, and 2020. Data for all years were downloaded from SEDAC in GeoTiff format at 2.5 min resolution (equivalent to ~5km grids), and converted to GPS coordinates using the R package 'raster' (21).

To estimate the population density for a given sampling location, we calculated the distance between the GPS coordinates for the sampling location and all ~5km grids in Kenya using the haversine distance formula as implemented in the R package 'geosphere' (22). We extracted the $\log_{10}$ 2020 population density estimate for the grid with the shortest distance to the sampling location and used this value as a potential mediating variable in downstream analyses.

To estimate the population density of each individual's birth location around the time they were born, we first assigned GPS coordinates to each birth location using Google Maps. We then found the ~5km grid closest to the birth location GPS coordinates using the procedure described above. Next, we estimated the population density of the grid during the year the individual was born, by fitting a linear model of $\log_{10}$ population density as function of year (using all available data, from 2000, 2005, 2010, 2015, and 2020). We extracted the beta and intercept from this linear model, and calculated $\log_{10}$ population density for the individual's birth year as intercept + beta x birth year.

*Distance to closest major city*

Population estimates in 2019 for major cities in Kenya were downloaded from the World Population Review (http://worldpopulationreview.com/countries/kenya-population/cities/). We considered major cities to be those with population counts >10,000, and extracted GPS locations for these cities from Google Maps. Using GPS coordinates for each sampling location, we calculated the distance between all major cities and the sampling location using the haversine distance formula as implemented in the R package 'geosphere' (22). We used the shortest distance (in kilometers) from each sampling location to a major city as potential mediating variable in downstream analyses.

**Identifying factors that mediate lifestyle effects on measured biomarkers**

For each biomarker that was significantly associated with lifestyle (Table S5), we were interested in identifying specific variables that mediated urban-rural differences health. However, we did not perform mediation analyses for lipid traits and for blood glucose, as sample sizes for these biomarkers were smaller to begin with, and after overlapping with our dietary data we could only include 50-60 urban individuals. Therefore, we focused mediation analyses on waist circumference, BMI, diastolic and systolic blood pressure, body fat, and our composite measure of health (sample sizes for mediation analyses are presented in Table S7).

To implement mediation analyses, we used an approach similar to (23, 24), to estimate the indirect effect of lifestyle on a given biomarker through the following potential mediating variables: alcohol and tobacco use (yes/no); consumption of meat, milk, blood, cooking oil, sugar, salt, bread, rice, ugali, potatoes, soda, fried foods, and sweets (frequency of use measured on a 0-4 scale); total number of unique carbohydrate items consumed; number of meals eaten per day; distance to the nearest city (in km); $\log_{10}$ population density; main subsistence activity; proportion of mothers in the sampling location with no formal education; and a tally of the number of market-derived amenities an individual possessed (as described in *Interview and food* *frequency questionnaire data*). Occupation was coded to reflect integration in the market economy as follows: 0 = animal keeping, farming, fishing, hunting and gathering; 1 = charcoal burning, mat making; 2 = casual worker, petty trade, self-employment; 3 = formal employment. To estimate the proportion of uneducated mothers in a given area, we calculated the fraction of women sampled in a given area with >0 children who reported having received no education. Population density, distance to a city, the proportion of uneducated mothers, and the number of owned market goods are all measures that have been used in the literature to describe how urban a given individual/location is (8, 9)

For all mediation analyses, we used two categories to describe lifestyle, urban and rural, given that previous analyses showed minimal health differences between pastoralist and non-pastoralist Turkana living in rural areas. For biomarkers with no sex x lifestyle interaction, we estimated the strength of the indirect effect of each mediator as the difference between the effect of lifestyle (urban versus rural) in two linear models: the 'unadjusted' model that did not account for the mediator (equivalent to model (1) in *Testing for lifestyle effects on measured biomarkers*), and the effect of lifestyle in an 'adjusted' model that also incorporated the mediator. If a given variable is a strong mediator, the effect of lifestyle will decrease when this variable is included in the model and absorbs variance otherwise attributed to lifestyle. For each biomarker, the adjusted model was implemented as follows:

$$y_i = \beta_0 + l_i\beta_l + a_i\beta_a + s_i\beta_s + m_i\beta_m + e_i \qquad (3)$$

Where $\beta_m$ represents the effect of the potential mediator on the outcome variable (all other variables are as defined in model (1)). For body fat percentage, which displayed lifestyle effects in females but not males (Figure S1), we modeled females only and removed the sex term ($\beta_s$)

from models (1) and (3). For the composite measure of health, we used generalized linear models with a binomial link function instead of linear models.

To assess the significance of each mediating variable, we estimated the decrease in $\beta_l$ in model (1) relative to model (3) across 1000 iterations of bootstrap resampling. We deemed a variable to be a significant mediator if the lower bound of the 95% confidence interval (for the decrease in $\beta_l$) did not overlap with 0. As a measure of effect size, we report the proportion of 1000 bootstrap resampling iterations for which the effect of lifestyle ($\beta_l$) was reduced when the potential mediating variable was included in the model (Table S7); a proportion >0.975 is equivalent to a 95% confidence interval that does not overlap with 0. As another measure of effect size, we report the percent change in the lifestyle effect estimated from model (1) relative to model (3) for each biomarker-mediator pair (without bootstrapping and using the full dataset to estimate each effect size; Figure 3). Further, to understand the degree to which the total set of mediators we identified explain the relationship between lifestyle and a given biomarker, we report the percent change in effect size for the lifestyle effect estimated from model (1) versus a model that included all the same covariates as well as all significant mediators for a given biomarker.

**Testing for early life effects on biomarkers of adult health**

Early life environments can have long-lasting effects on health, such that conditions experienced during the first years of life may predict health outcomes measured decades later in adulthood. In humans, associations between early life resource constraint and adult health and disease are often interpreted as 'predictive adaptive responses' (PAR) gone wrong. Specifically, it is hypothesized that individuals experiencing early life resource scarcity develop phenotypes well-suited to scarcity; if they instead encounter resource abundance later in life, health problems ensue (25, 26). A second alternative hypothesis, commonly referred to as the ‘developmental constraints’ or ‘silver spoon’ hypothesis, posits that associations between early life challenges and compromised adult health are the result of simple costs to health incurred from a suboptimal developmental environment (27–29). Importantly, these two hypotheses (PAR and developmental constraints) make contrasting predictions. While PAR predicts that individuals experiencing matched early life and adult conditions should experience better health than individuals experiencing mismatched environments across the life course, developmental constraints predicts simple main effects of early life conditions (regardless of the adult environment). Several research groups (30–34) have operationalized these hypotheses by asking whether there is evidence for crossing reaction norms/interaction effects between early life and adult environments (in support of PAR) or whether early life adversity is simply associated with compromised adult health (in support of developmental constraints). We took a similar approach to disentangle these hypotheses.

For biomarkers with no sex x lifestyle interaction, we first asked whether there was any evidence for interaction effects between adult lifestyle and lifestyle/urbanicity during early life using the following linear model:

$$y_i = \beta_0 + l_i\beta_l + a_i\beta_a + s_i\beta_s + d_i\beta_d + (l_i * d_i)\beta_{l*d} + e_i \qquad (4)$$

Where $d_i$ represents the $\log_{10}$ population density for the birth location of individual $i$ during their birth year, $l_i$ represents adult lifestyle (urban versus rural), and $\beta_{l*d}$ captures the interaction effect between these two variables. For the two variables with lifestyle effects on health in females but not males (body fat percentage and blood glucose levels), we modeled females only and removed the sex term ($\beta_s$). For the composite measure of health, we used generalized linear models with a binomial link function instead of linear models. For each of the 11 models, we extracted the p-value associated with $\beta_{l*d}$ and corrected for multiple hypothesis testing (12). In all cases, the nominal and FDR-corrected p-value was >0.05, suggesting predictive adaptive responses do not explain early life effects on health in the Turkana.

Next, we reran the appropriate version of model (4) for each measure after removing the interaction effect ($\beta_{l*d}$). For biomarkers with no sex x lifestyle interaction, this model was equivalent to:

$$y_i = \beta_0 + l_i\beta_l + a_i\beta_a + s_i\beta_s + d_i\beta_d + e_i \tag{5}$$

For each of the 11 models, we extracted the p-value associated with early life effect ($\beta_d$) and corrected for multiple hypothesis testing (12). We considered a given variable to show support for the developmental constraints hypothesis if the FDR-corrected p-value was less than 0.05. Results for all models described in this section are presented in Table S8. Results for parallel analyses that use $\log_{10}$ population density for the sampling location to define the adult environment (rather than a binary urban/rural lifestyle variable) are presented in Table S9.

**Translating estimates of effect size into percentages**

In the main text, we report several estimates of effect size as the percent difference between, for example, urban and rural individuals, to help the reader contextualize the magnitude of biomarker differences we observe. To obtain these percentages, we translated our fitted linear mixed effects model from model (1) or model (5) into percent differences in each biomarker as a function of the early life or adult environment.

To estimate the percent difference in, for example, BMI between an individual living in a rural versus urban location, we reran model 1 without normalizing the outcome variable. We then extracted the fitted model estimates (for $\beta_0, \beta_l, \beta_a, \beta_s$) and used them to calculate BMI for an 'average' individual in the population by summing the fitted effect sizes multiplied by average values from the dataset ($a_i$=40.4 years; $s_i$=0 (indicating female)). To isolate how lifestyle influenced BMI, we performed these calculations after setting $l_i = 1$ (indicating rural, Turkana county) or $l_i = 0$ (indicating urban, Laikipia county). Finally, we calculated the percent difference between the two BMI estimates, which reflect values for an average female living in each location.

We performed parallel calculations using the fitted model estimates from model (5), in order to understand the relative impact of early life versus adult environmental conditions. Here, we extracted the estimates for $\beta_0, \beta_l, \beta_a, \beta_s, \beta_d$, and varied $d_i$ or $l_i$ to isolate the effects of early life population density or adult lifestyle, respectively. For calculations involving early life population density, $l_i$ was set to 1, and $d_i$ was varied between 0.504 and 1.360, which represent the 25$^{th}$ and 75$^{th}$ percentiles of $\log_{10}$ early life population densities. We calculated percent differences in health biomarkers in this manner for the biomarkers with significant early life effects as estimated in model (5), namely: average waist circumference, BMI, diastolic blood pressure, body fat percentage, and the composite measure of health (Table S8). For body fat percentage, where environmental effects are only observed in females, all models focused on this sex alone and did not include the effect of $\beta_s$.

### Supplementary Figures

**Figure S1. Mean biomarker values for measures with a significant sex x lifestyle interaction.** Using likelihood ratio tests, we determined that (A) body fat and (B) blood glucose had significant sex x lifestyle interactions (Table S5), such that lifestyle effects were observed in females, but not males. Raw data are shown to support this conclusion, all contrasts between the three lifestyle categories analyzed for males alone were not significant (linear model controlling for age, all $p>0.05$, sample sizes as in Table S1). Contrasts between the three lifestyle categories analyzed for females alone showed a similar pattern to what is described in the main text for other biomarkers (Figure 2). Specifically, no significant differences were observed between body fat ($p=0.391$) and blood glucose levels ($p=0.629$) for pastoralist versus rural, non-pastoralist Turkana; however, strong differences were observed between both of these groups and urban, non-pastoralist Turkana (contrast between pastoralists and urban non-pastoralists: body fat, $p=4.660\times10^{-5}$, blood glucose: $p=2.179\times10^{-7}$; contrast between rural and urban non-pastoralists: body fat, $p=3.194\times10^{-5}$, blood glucose: $p=4.030\times10^{-4}$).

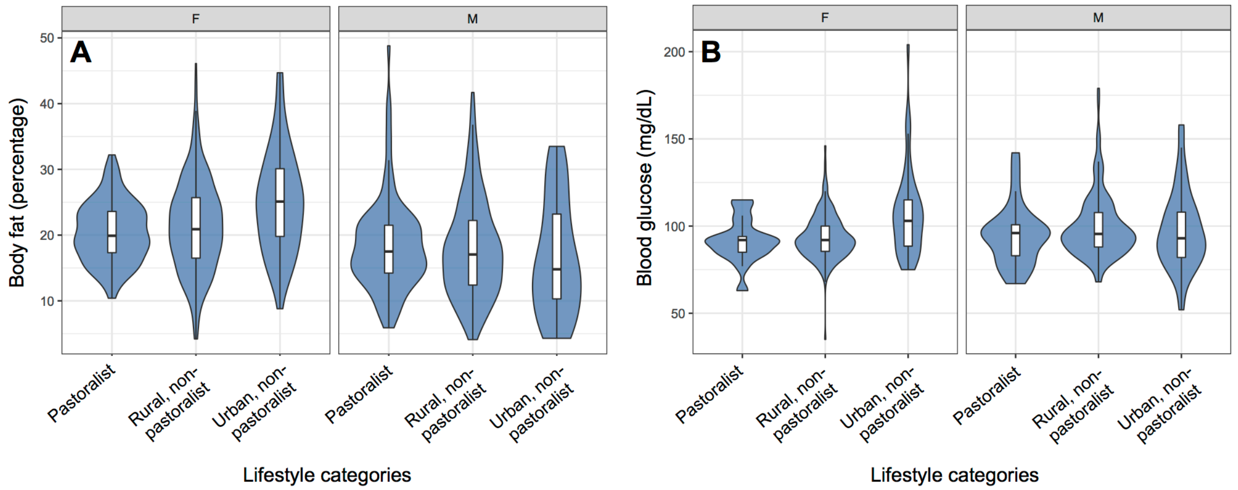

**Figure S2. Mean biomarker values for Turkana individuals versus the US (NHANES).** Raw data for 8 measured biomarkers are shown for the NHANES dataset (filtered to include individuals aged >18 and <65 years old) versus rural and urban Turkana. Raw data for the two blood pressure measures are shown in Figure S4, and data for blood glucose are not shown as no significant differences between groups were found (Figure 2 and Table S5). Outliers in the most extreme 1% of the dataset for BMI, total cholesterol, triglycerides, and waist circumference are not shown to aid visualization. For body fat, where only females were analyzed, only data for this sex is plotted. Units of measure are as in Table S1.

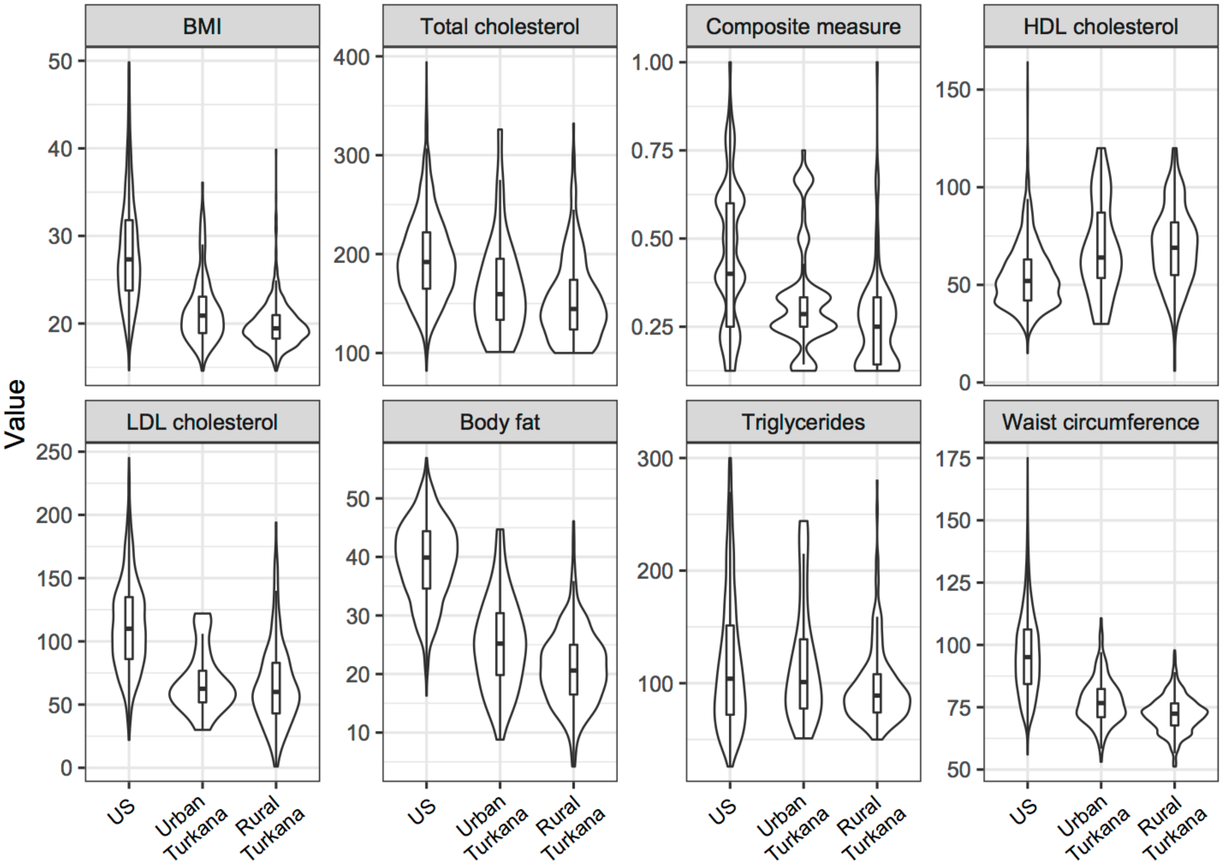

**Figure S3. Proportion of Turkana versus US (NHANES) individuals whose biomarker values exceed clinical cutoffs.** Cutoffs were defined as follows: (i) waist circumference >89 cm for women or >102 cm for men; (ii) triglyceride levels >150 mg/dL; (iii) HDL cholesterol levels <40 mg/dL for men or <50 mg/dL for women; (iv) blood pressure >135/85; (v) BMI >25; (vi) LDL cholesterol levels >100 mg/dL; and (vii) total cholesterol levels > 200 mg/dL. Cutoffs for BMI, LDL cholesterol, and total cholesterol were taken from CDC recommendations. Cutoffs for the remaining biomarkers followed the criteria used by the American Heart Association to define metabolic syndrome (11). For blood pressure, we also show hypertension prevalence in the US when we expand our definition to include individuals taking medications used to treat high blood pressure (14.85% of US individuals, shown as open bar extension); no Turkana reported being on high blood pressure medications. Only biomarkers that exhibited significant differences between the US and Turkana are shown.

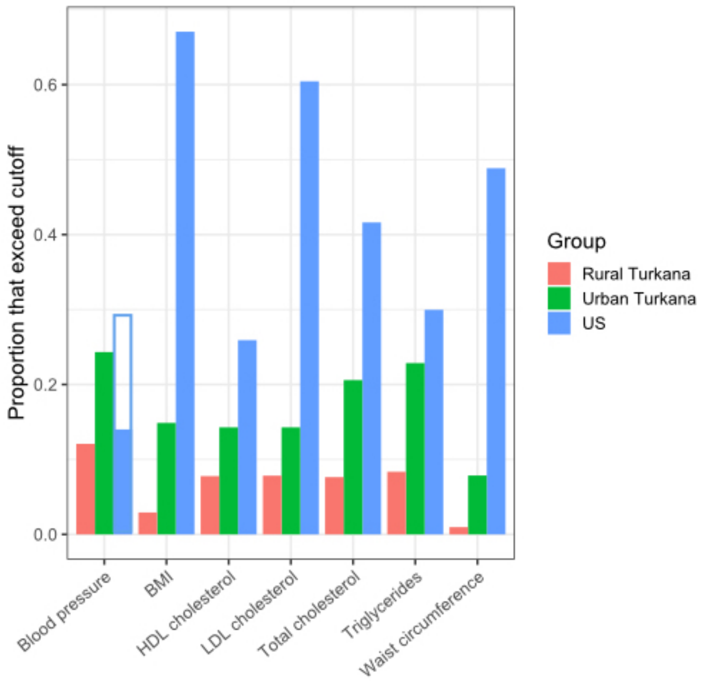

**Figure S4. Blood pressure values for Turkana individuals versus the US (NHANES).** Raw data for (A) diastolic and (B) systolic blood pressure are shown for the NHANES dataset (filtered to include individuals aged >18 and <65 years old) versus rural and urban Turkana. Outliers in the most extreme 1% of the dataset are not shown to aid visualization. Loess curves against age are shown for each population.

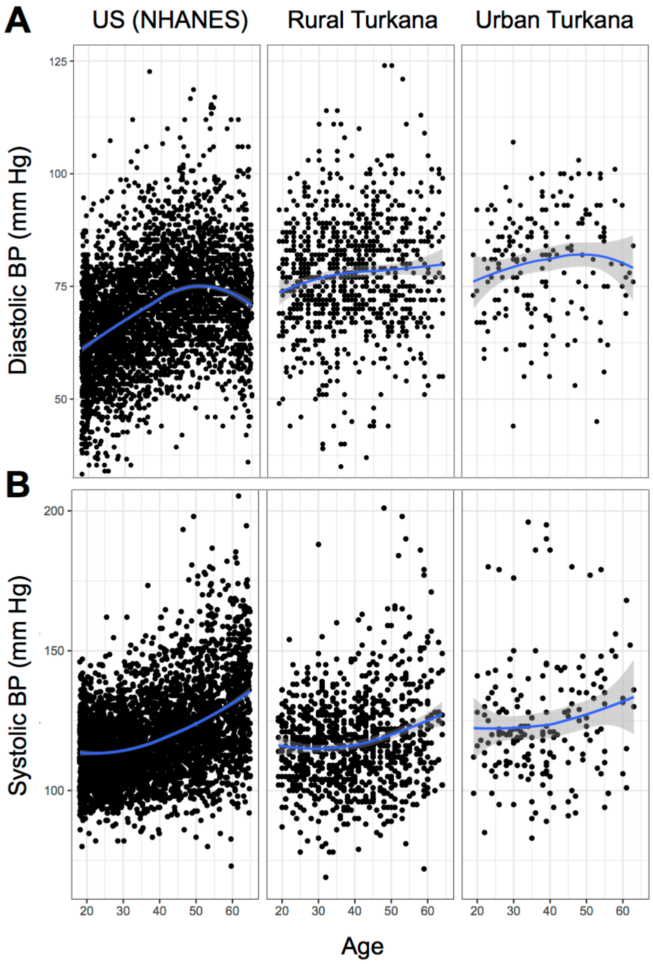

**Figure S5. Relationship between birth location and location at time of sampling.** (A) Red points denote birth locations of study participants, open circles denote sampling locations, and grey lines connect each participant's birthplace with the location at which they were sampled in adulthood. No lines are drawn for participants that were born and sampled in the same location. (B) Distance between birth location and sampling location for participants sampled in urban, Laikipia county versus participants sampled in rural, Turkana county. Participants sampled in Laikipia county are more likely to be living far from their birth location (Wilcoxon test, $p=1.364 \times 10^{-15}$). (C) There is minimal correlation between the population density of each individual's birth and sampling locations, suggesting that effects of urbanicity across the life course can be considered independently ($R^2 = 0.115$, $p<10^{-16}$).

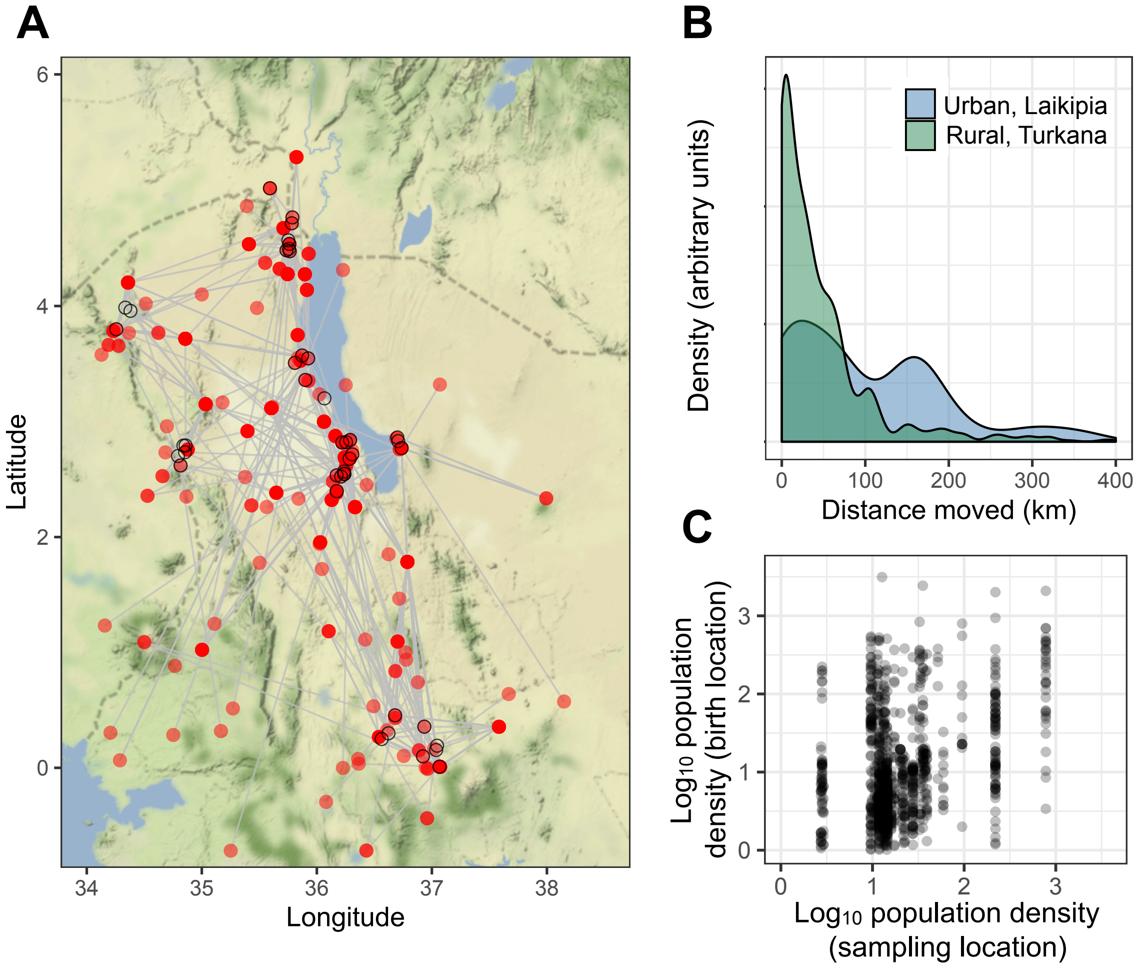

**Figure S6. Correlation coefficients among measured biomarkers.** Matrix shows the pairwise correlation for each pair of measured biomarkers, after controlling for age and sex. Specifically, we regressed out age and sex and estimated the Pearson correlation coefficient between each pair of residuals. Only significant ($p<0.05$) correlations are shown, and are visualized using the 'corrplot' package in R (35). The strongest correlation we observed is between total and LDL cholesterol levels ($R^2$=0.762), all other biomarker pairs are only weakly to moderately correlated ($R^2$<0.35). We note that lifestyle effects were found for total cholesterol, but not LDL cholesterol levels, suggesting that collinearity between these two measures did not produce false positive results.

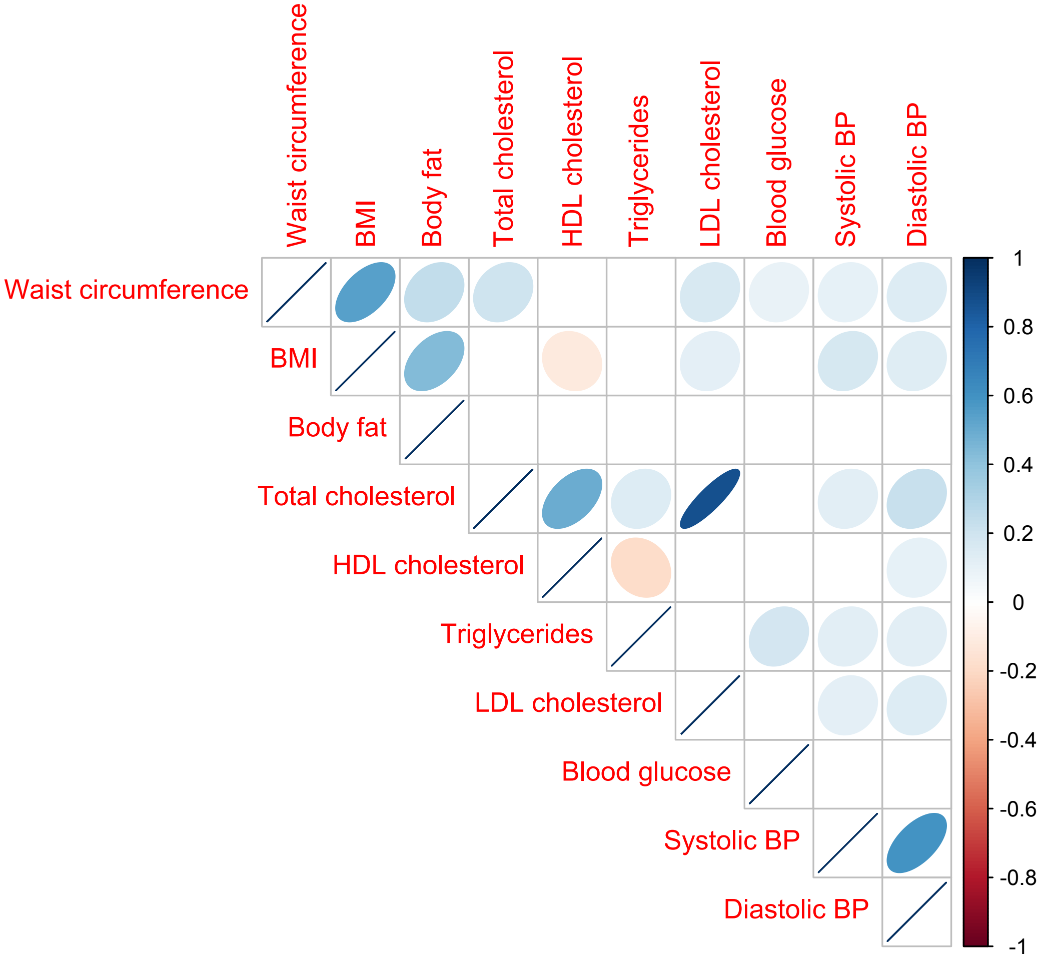
